# Label-Free Composition Analysis of Supramolecular Polymer – Nanoparticle Hydrogels by Reversed-Phase Liquid Chromatography Coupled with a Charged Aerosol Detector

**DOI:** 10.1101/2023.08.11.553055

**Authors:** Shijia Tang, Zachary Pederson, Emily L. Meany, Chun-Wan Yen, Andrew K. Swansiger, James S. Prell, Bifan Chen, Abigail K. Grosskopf, Noah Eckman, Grace Jiang, Julie Baillet, Jackson D. Pellett, Eric A. Appel

**Affiliations:** Synthetic Molecule Pharmaceutical Sciences, Genentech, Inc., South San Francisco, CA 94080, USA; Department of Bioengineering, Stanford University, Stanford, CA 94305, USA; Department of Chemical Engineering, Stanford University, Stanford, CA 94305, USA; Department of Materials Science and Engineering, Stanford University, Stanford, CA 94305, USA; Department of Chemistry and Biochemistry, University of Oregon, Eugene, Oregon 97403, USA; Preclinical and Translational Pharmacokinetics and Pharmacodynamics, Genentech, Inc., South San Francisco, CA 94080, USA

**Keywords:** Charged aerosol detection, polymer quantification, reversed-phase separation, supramolecular, hydrogel

## Abstract

Supramolecular hydrogels formed through polymer-nanoparticle interactions are promising biocompatible materials for translational medicines. This class of hydrogels exhibits shear-thinning behavior and rapid recovery of mechanical properties following applied stresses, providing desirable attributes for formulating sprayable and injectable therapeutics. Characterization of hydrogel composition and loading of encapsulated drugs is critical to achieving desired rheological behavior as well as tunable in vitro and in vivo payload release kinetics. However, quantitation of hydrogel compositions is challenging due to material complexity, heterogeneity, high molecular weight, and the lack of chromophores. Here, we present a label-free approach to simultaneously determine hydrogel polymeric components and encapsulated payloads by coupling a reversed phase liquid chromatographic method with a charged aerosol detector (RPLC-CAD). The hydrogel studied consists of modified hydroxypropylmethylcellulose, self-assembled PEG-b-PLA nanoparticles, and a therapeutic compound, Bimatoprost. The three components were resolved and quantitated using the RPLC-CAD method with a C4 stationary phase. The method demonstrated robust performance, applicability to alternative cargos (i.e. proteins), and was suitable for composition analysis as well as for evaluating in vitro release of cargos from the hydrogel. Moreover, this method can be used to monitor polymer degradation and material stability, which can be further elucidated by coupling the RPLC method with high resolution mass spectrometry and a Fourier-transform based deconvolution algorithm. To our knowledge, this is the first RPLC-CAD method for characterizing the critical quality attributes of supramolecular hydrogels. We envision this analytical strategy could be generalized to characterize other classes of supramolecular hydrogels, establish structure-property relationships, and provide rational design guidance in hydrogel drug product development.

## 1. Introduction

Supramolecular hydrogels are physically cross-linked viscoelastic biomaterials that are rapidly expanding in drug delivery, cell therapy, surgical coatings, medical device applications, and beyond.[1–10] Through tuning the chemistries and crosslinking density (mesh size of a hydrogel molecular network), hydrogels can be made that adopt vastly different chemical or physical properties and can encapsulate a variety of cargoes and accommodate different targeted release time frames.[3, 6–7] In comparison to chemically cross-linked hydrogels, supramolecular hydrogels rely on physical, non-covalent interactions such as ionic interactions, hydrophobic interactions, hydrogen-bonding, metal-ligand complexation, host-guest complexation, or biorecognition, which provide several clinical and process development benefits, such as gelation without reactive moieties or volume change.[1, 6, 8] Moreover, the reversible, non-covalent interactions in supramolecular hydrogels form dynamic and transient crosslinks, resulting in rapid self-healing and shear-thinning properties that make these hydrogels an ideal formulation strategy for sprayable and injectable therapeutics.[6–7]

While the materials library of supramolecular hydrogels is expanding, few analytical methods have been developed to quantitatively characterize the hydrogel components. Currently, published studies mainly focus on the content and release of encapsulated drugs. The quantitation of hydrogel matrix polymers, in addition to drug payloads, is important to establish structure-property relationships and provide information on critical quality attributes (CQAs). For example, comparing polymer and drug release profiles can shed light on the release mechanisms (i.e., driven by diffusion, erosion), pharmacokinetics, fate of the matrix polymers over time, and establish in vitro and in vivo correlation (IVIVC), and thereby enabling the rational design of hydrogels for specific target product profiles.[3]

Several challenges are inherent to the compositional analysis of supramolecular hydrogels. From a chromatography perspective, hydrogels often contain both encapsulated payloads and matrix, typically consisting of two or more high molecular weight and heterogeneous polymeric components. This requires a method that resolves multiple components while allowing good recovery for the polymers. In addition, an appropriate sample preparation procedure is critical to dissociate the hydrogels and fully extract the individual components without degradation. Due to these challenges, hydrogel polymeric components have typically been characterized only at the precursor stage rather than after hydrogel formation. In the final hydrogel product, only the active payload is typically quantitated to determine the loading or release profiles.[11–14] From a detection perspective, the encapsulated payloads are often UV active, while many polymers lack UV chromophores and require derivatization or an alternative detection principle to quantify. Labeling approaches, such as modifying the polymeric components with fluorescent tags or encapsulating fluorescent dyes as the payload surrogate, have been previously developed for tracking polymer and payload release.[15–20] However, labeling approaches can complicate hydrogel chemistries and/or release kinetics depending on the degree of modification and fluorescent modifier properties. Additionally, tracking fluorescence intensity may not fully reflect any chemical changes in the polymer backbones over time. Identifying a label-free approach that combines chromatography separation with a universal detection technique for non-UV absorbing compounds would be beneficial to realize quantitation for all components in a supramolecular hydrogel and capture key chemical changes over time. However, the label-free hydrogel analysis is rarely explored in the literature, and it remains a gap on what chromatographic separation modes and detection techniques can provide sufficient sensitivity, resolution, and recovery for all the components undergoing quantitative analysis.

Recently, a supramolecular hydrogel platform employing polymer-nanoparticle interactions between dodecyl-modified hydroxypropylmethylcellulose (HPMC-C_12_) and poly(ethylene glycol)-block-poly(lactic acid) nanoparticles (PEG-b-PLA NPs) has been developed, which demonstrates exceptional injectability and rapid self-healing properties (**Scheme 1**).[21–26] These materials are denoted as PNP-X-Y, where X refers to the weight percent loading of the HPMC-C_12_ component and Y refers to the weight percent loading of the PEG-b-PLA NP component (e.g., PNP-2-10 gels comprise 2 wt% HPMC-C_12_ and 10 wt% PEG-b-PLA NPs). In this study, we use PNP-2-10 hydrogel as a model system to develop a label-free analytical method utilizing reversed phase liquid chromatography coupled to a charged aerosol detector (RPLC-CAD) which quantitates all components in the hydrogel—HPMC-C_12_, PEG-b-PLA NPs, and encapsulated therapeutic payload Bimatoprost. A wide pore reversed phase column was selected and provided great specificity and recovery for all the hydrogel components, whereas size exclusion chromatography (SEC) did not resolve the HPMC-C_12_ and the PEG-b-PLA NPs. Due to the lack of UV chromophores on both polymeric components, a highly sensitive aerosol-based detection technique, CAD, was identified as most suitable to couple with the RPLC separation for quantitative analysis instead of differential refractometer or light scattering techniques. Our developed method demonstrated a label-free approach that provided high specificity, precision, accuracy, and sensitivity, allowing for determining CQAs for hydrogels, i.e., simultaneous quantitation of hydrogel polymeric components and drug cargos. The method was capable of differentiating polymer integrity after degradation or E-beam sterilization and could be combined with mass spectrometry (MS) for further structural elucidation and monitoring of material stability. The method was also applicable to an alternative cargo, Bovine Serum Albumin (BSA), and demonstrated that this methodology can be generalized to characterize other supramolecular hydrogels with various modalities of payloads to gain more chemical and process understanding of these highly complex biomaterials and progress their clinical translation.

**Scheme 1.**
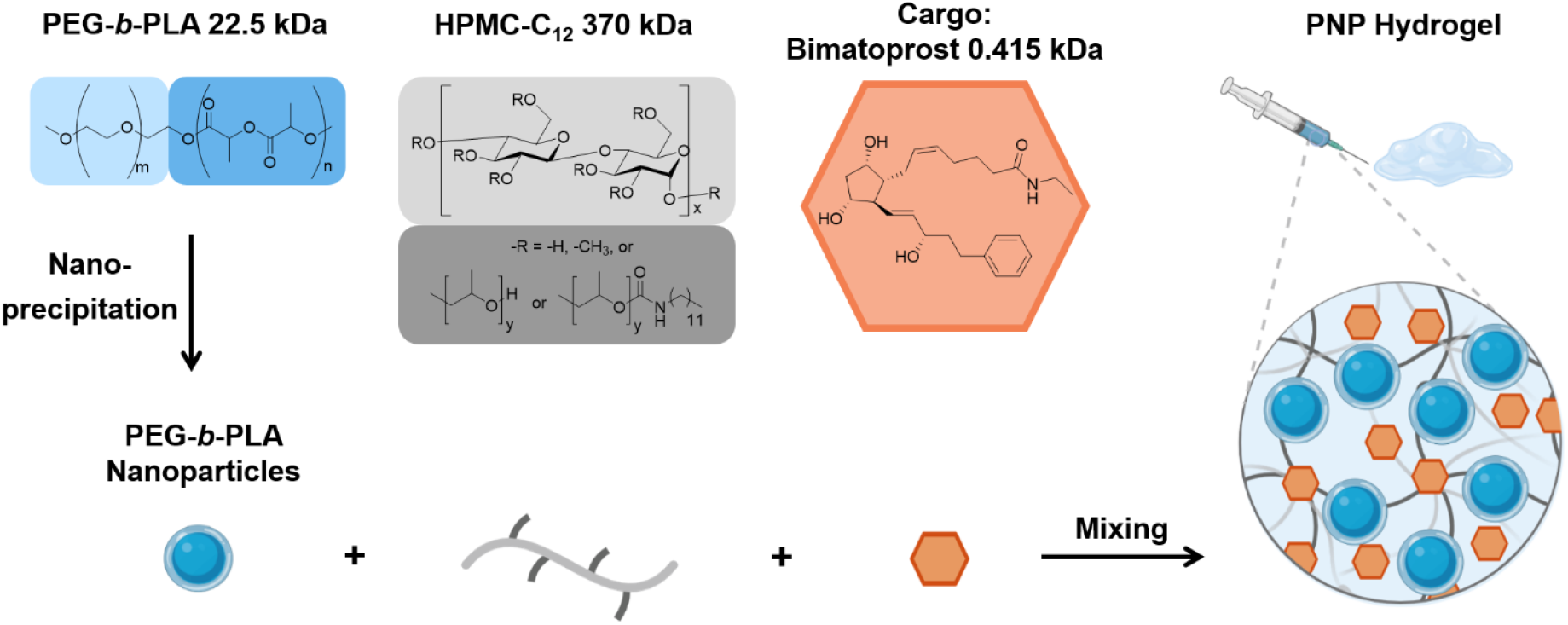
Supramolecular Polymer-Nanoparticle Hydrogel Composition and Gelation Process

## 2. Results and Discussion

### 2.1. Separation of Hydrogel Components by SEC and RPLC

The supramolecular PNP hydrogel contains components with vastly different molecular weights, conformation, and hydrophobicity (Scheme 1): the encapsulated small molecule cargo Bimatoprost and hydrogel matrix components HPMC-C_12_ and polymeric nanoparticles formed by nanoprecipitation of PEG-b-PLA copolymers. We first focused on identifying a separation mode that could provide good resolving power and recovery for all three components. SEC is the benchmark method for polymer analysis, which separates analytes based on their hydrodynamic radius, Rh.[32] Three SEC columns (Acclaim SEC-1000, PolySep GFC-P6000, and TOSOH TSKgel G5000PWXL) composed of hydrophilic polymer beads designed for the separation of high Mw water soluble polymers were assessed by connecting to a CAD (**Table S1**, Supporting Information). For all the SEC-CAD methods, ammonium acetate was used as the MP modifier for compatibility with CAD and 10-20% (v/v) MeCN was required to promote the elution of the PEG-b-PLA NPs, likely due to the secondary interaction between the analytes and the column phases. **Figure 1(a)** displays a representative SEC-CAD chromatogram obtained using the TOSOH TSKgel G5000PWXL. A sufficient resolution could be achieved among the four pullulan sizing standards (1330 kDa to 0.99 kDa) in the Mw range of the hydrogel polymers. However, HPMC-C_12_ and PEG-b-PLA NPs showed co-elution in the SEC (Figure 1(a)). We further investigated the co-elution by coupling SEC to MALS and an inline viscometer (IV) to determine Mw and Rh. The IV analysis revealed the HPMC-C_12_ and the PEG-b-PLA NPs had similar Rh (**Table S2**, Supporting Information), which combined with the apparent high dispersity of HPMC-C_12_ (peak width ∼10 min at baseline), suggested the resolving power of SEC was insufficient in the hydrogel analysis.

**Figure 1.**
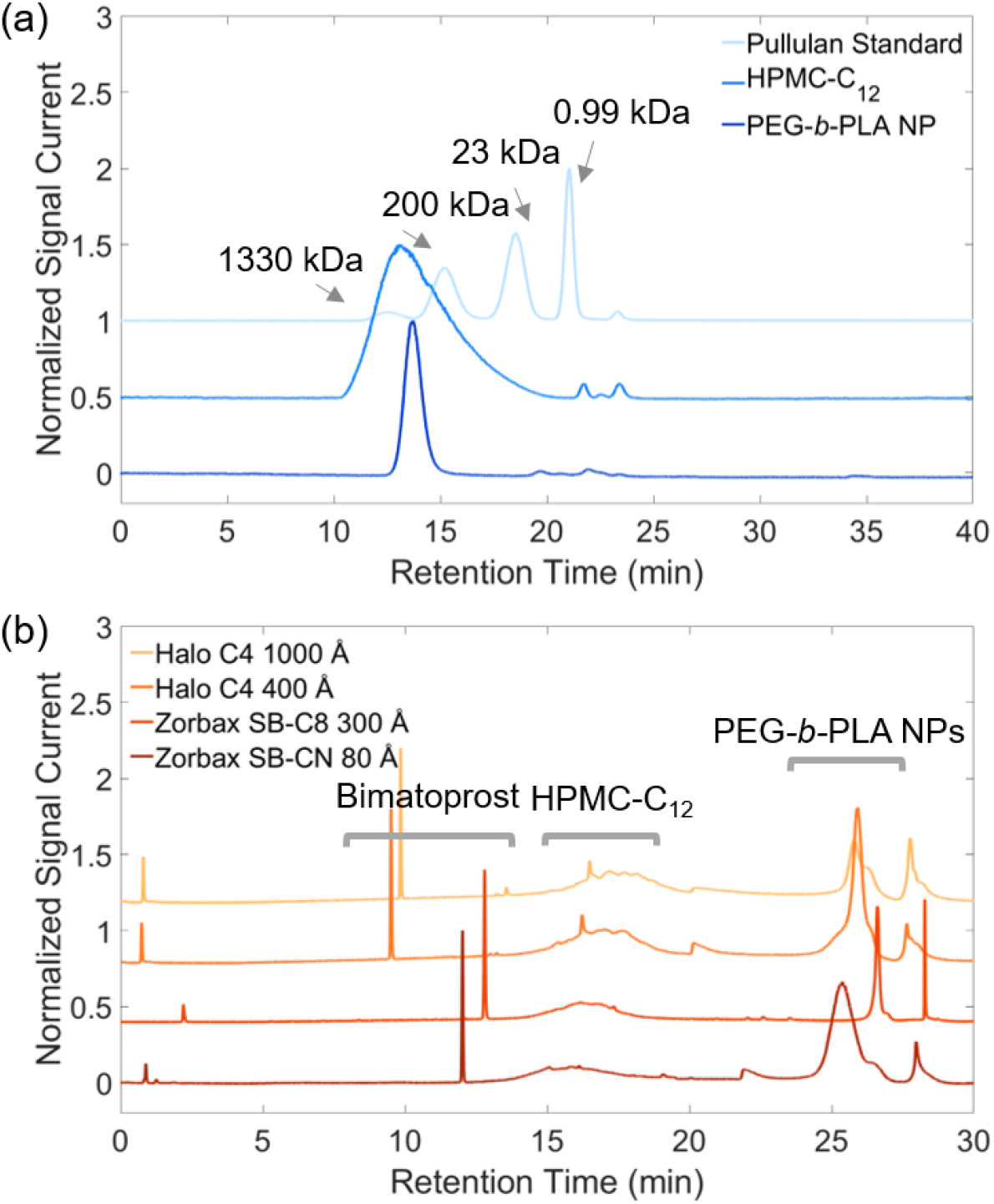
Representative chromatograms of Bimatoprost, HPMC-C12, and PEG-b-PLA NPs using different separation principles: **(a) SEC-CAD and (b) RPLC-CAD.** Given the insufficient resolving power of the size based method, reversed-phase separation was selected that can potentially provide better selectivity for HPMC-C_12_ and PEG-b-PLA NPs. However, it is a less common separation mode used in nanoparticle characterization limited by the small pore size of the column packing materials. RP columns with 80-1000 Å pore size and less hydrophobic stationary phase chemistries were selected to allow the elution of HPMC-C_12_ and PEG-b-PLA NPs (Table S1, Supporting Information). Figure 1(b) shows a comparison of four RP columns using a generic linear gradient with a thermostat temperature of 30 °C. All RP columns showed improved resolution between the 3 analytes. In the Zorbax SB-CN analysis (Figure 1(b)), both HPMC-C_12_ and PEG-b-PLA NPs peaks were broad, likely due to restricted diffusion of the NPs since the column is packed with fully porous particles (FPP) with a pore size of 80 Å.[33] The peak shape of HPMC-C_12_ and PEG-b-PLA NPs were improved by switching to a 300 Å C8 FPP column (Figure 1(b)), although the retention of the PEG-b-PLA NPs was quite strong by the hydrophobic stationary phases. To further improve the mass transfer kinetics and elution, Halo C4 400 Å and 1000 Å columns packed with superficially porous particles (SPP) and less hydrophobic phases were tested. Both columns showed improved efficiency in HPMC-C_12_, better resolution between Bimatoprost and HPMC-C_12_ compared to the C8 column, while providing reasonable peak shapes for the PEG-b-PLA NPs. Therefore, the C4 columns were pursued for further optimizations.

Also notably, driven by the hydrophobic interaction of the PLA segments, the PEG-b-PLA can spontaneously form aggregates on column when exposed to an aqueous environment (i.e., in a chromatographic diluent or an HPLC system). The IV analysis showed that dissolving the PEG-b-PLA copolymers in an HPLC diluent (75/25 H2O/MeCN, v/v), designated as the PEG-b-PLA standard (PEG-b-PLA STD), led to the Mw increasing from 22.5 kDa as a single copolymer chain to ∼12 MDa with a hydrodynamic radius of 17.2 nm, indicating the formation of nanosize aggregates on column (Table S2, Supporting Information). The Rh of the nanoparticles in PEG-b-PLA STD was comparable to the ones in PEG-b-PLA NPs formed by nanoprecipitation (Rh = 25.4 nm, Table S2, Supporting Information), with PDIPEG-b-PLA STD being 1.2 and PDIPEG-b-PLA NP being 1.0. The higher PDI in PEG-b-PLA STD was consistent with the self-assembly process of the copolymers without homogenization or filtering, leading to more heterogeneous size distribution in the aggregates. Nevertheless, the comparable Rh of the PEG-b-PLA STD and the PEG-b-PLA NPs allowed the quantitation of PEG-b-PLA NPs in their native spherical conformation using the aggregates in PEG-b-PLA STD as the external standard.

### 2.2 Hydrogel development and characterization

Recovery and Sensitivity Optimization: Our next focus was to reduce the PEG-b-PLA NPs carryover and improve the HPMC-C_12_ peak shape and height to achieve better sensitivity. Counterintuitively, increasing the column pore size from 400 to 1000 Å led to a higher PEG-b-PLA NPs carryover at 30 °C (**Table 1**). The 400 Å C4 column showed a 6.2% carryover compared to the 1000 Å C4 column at 23.0%. Considering the size of the PEG-b-PLA NPs and the starting gradient of 5% MeCN was not a thermodynamically good solvent for the NPs, the NPs were more likely to enter, precipitate and partition into the 1000 Å pores – operating in an interaction/adsorption mode and led to more carryover.[34] In contrast, the 400 Å C4 column likely operated with a hybrid mode of exclusion (entropic) and interaction (enthalpic), considering the pore size is close to the NPs Rh. When the thermostat temperature was elevated to 50 °C, a reduction in the mobile phase viscosity and an increase in the analyte diffusivity accelerated mass transfer and absorption/desorption, shifting the interaction dominant separation to exclusion dominant separation, and thereby reducing the carryover in both 1000 Å and 400 Å (< 2% above 50 °C).[34–35] Considering the 400 Å column had less tendency to trap PEG-b-PLA NPs in the pores, this column was selected in the final method.

**Table 1.** Effect of Separation Temperature on the Carryover% of PEG-b-PLA.

| Pore Size | Carryover (% Area/Area)<sup>a</sup> |  |  |  |
| --- | --- | --- | --- | --- |
|  | 30 °C | 40 °C | 50 °C | 60 °C |
| 1000 Å | 23.0% | 7.2% | <2% | <2% |
| 400 Å | 6.2% | 6.4% | <2% | <2% |

<sup>a</sup> Carryover% was determined by the ratio of the PEG-*b*-PLA peak area in the subsequent blank injection and the preceding PEG-*b*-PLA standard injection.

To further improve the peak shape and height for HPMC-C_12_, the thermostat temperature was increased to 50 °C. However, it did not improve the peak shape significantly and led to an increase in retention (**Figure S1(b),** Supporting Information). This can be explained by the temperature dependent gelation of HPMC. As temperature increases, the HPMC starts to lose the water shell, accompanied by an increase in the polymer-polymer interaction or polymer-stationary phase interaction.[36] A previous study reported gelation started at ∼26 °C, and the onset temperature of gelation can vary depending on the composition and the functionalization of HPMC.[37] To mitigate the impact of on-column gelation on the separation, while maintaining good recovery for PEG-b-PLA at 50-60 °C, a step-gradient was implemented to elute the HPMC-C_12_ and improve its on-column solubility (∼80% MeCN) (**Figure S1(c)**, Supporting Information). The sharpened HPMC-C_12_ peak suggested the on-column absorption had been alleviated and resulted in a fast elution. The sensitivity of the HPMC-C_12_ improved ∼5 fold compared to the initial gradient program. The cargo Bimatoprost was also assessed with the updated RPLC-CAD method, where peak splitting was observed, likely due to diluent incompatibility—50% MeCN diluent for Bimatoprost was too strong for the initial condition in the final method (10% MeCN). Therefore, using a weaker diluent was selected in order to quantitate all three components together and will be further discussed in the Diluent Study section.

### 2.3 Diluent Study

The diluent extraction efficiency of HPMC-C_12_ and PEG-b-PLA NPs was assessed to select a diluent and sample preparation protocol for quantitative analysis of the intact PNP-2-10 hydrogel. The extraction efficiency was determined by comparing the calculated amount of polymers from the calibration curve to the theoretical amount of polymers from the hydrogel label claim. Both a one-step dilution (gel dissolved as-is in the diluent) and a two-step dilution (gel dissolved in the organic portion first followed by adding the aqueous portion) were assessed (Figure 2). Organic solvent was essential to effectively disrupt the hydrophobic interaction between HPMC-C_12_ and PEG-b-PLA NPs. In an aqueous only diluent, the extraction efficiency for both HPMC-C_12_ and PEG-b-PLA NPs was poor (lower than 30%). The one-step and two-step dilution were performed with MeCN/H2O (25%/75%, v/v) instead of MeCN/H2O (50%/50%, v/v) due to the peak splitting observed for Bimatoprost with the latter diluent. The two-step dilution was more reliable compared to the one-step dilution. The one-step dilution showed more variations between duplicate preparations (data not shown). Two other organic solvents THF/H2O (50%/50%, v/v) and DMSO/H2O (50%/50%, v/v) were also assessed in the two-step preparation procedure, considering they have good solubility for the PEG-b-PLA NPs. However, DMSO showed poor extraction for both components. Although THF showed better extraction compared to DMSO, ultimately, the 2-step diluent MeCN/H2O (25%/75%, v/v) was selected as the final procedure considering (1) this diluent had good extraction efficiency reaching 90-110% of label claim for both polymeric components; and (2) the diluent strength was appropriate to avoid solvent incompatibility that caused peak splitting for hydrophilic cargo Bimatoprost.

**Figure 2.**
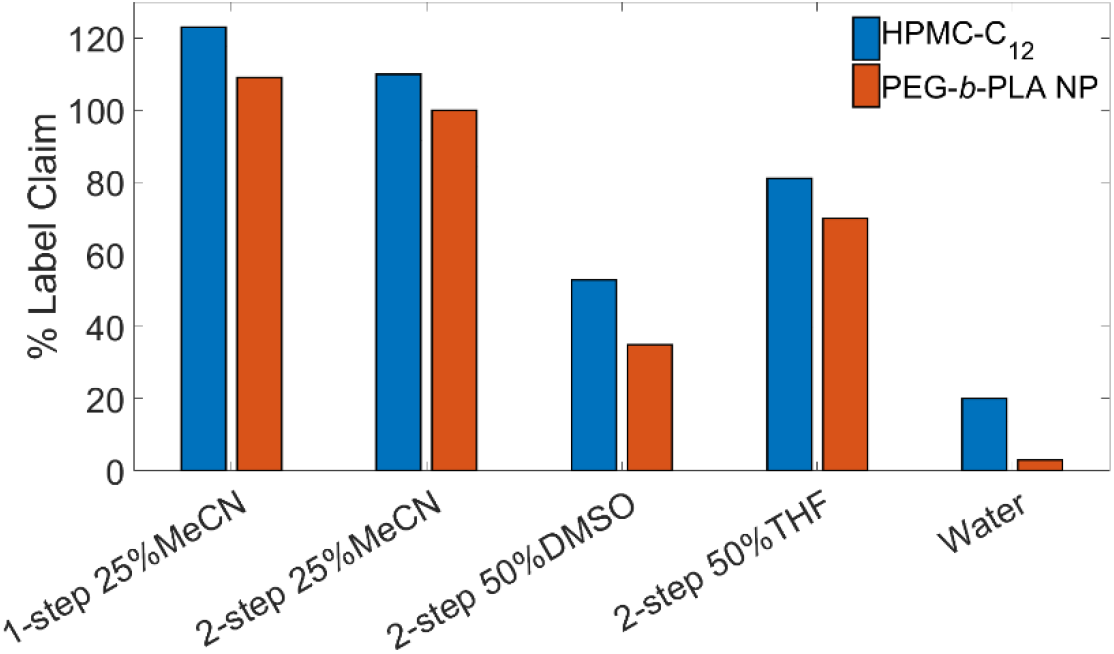
Effect of diluents and sample preparation on HPMC-C_12_ and PEG-b-PLA NPs extraction efficiency (n=2).

### 2.4 Method Performance

The final RPLC-CAD method condition was set at 60 °C to reduce carryover for PEG-b-PLA with the step-gradient for fast HPMC-C_12_ elution. The method performance was validated for specificity, linearity, precision, accuracy, and quantitation limit following ICH Q2 guidance (Figure 3). Specificity was shown by comparing the chromatograms of the PNP-2-10 hydrogel with the diluent blank (Figure 3(a)). No interfering peaks in the diluent blank were observed to affect the quantitation of all three components. The method was stability indicating, demonstrated through a forced degradation study by treating the HPMC-C_12_ (0.06 mg/mL) and PEG-b-PLA (0.1 mg/mL) standard mixture with acid (0.1 M HCl), base (0.1 M NaOH), or heat (60 °C) stressed conditions for ∼20 hr (Figure 3(b)). A common degradant was observed eluting ∼6 min (before the HPMC-C_12_), and it was formed most rapidly in the base stressed condition accompanied with loss of the PEG-b-PLA peak. The degradant was likely associated with the remaining PEG blocks after PLA blocks hydrolyzed and degraded (See Polymer Degradant Characterization by RPLC-MS Section). The method precision was determined by the %RSD of three replicate injections at 10%, 100% and 120% of the nominal sample loading. Each set of replicates have a %RSD lower than 3.0%, suggesting excellent method precision (Figure 3(c)). The accuracy of the method was within 90-110% (Figure 3(c), grey band) for all components at 10, 100, and 120% of the nominal loading level. The highest variation was observed for PEG-b-PLA, which can be explained by aggregates size variations in each PEG-b-PLA STD solution and the CAD response factor dependence on analyte size and molar mass.[38–39] Nevertheless, the response variation was acceptable and aligned with the inter-analyte CAD response factor’s variation in the previous study.[40] Finally, since one application of this CAD method was to study the in vitro release of the encapsulated cargo and matrix polymers, the method’s working range was validated spanning three orders of magnitude for each component in the PNP-2-10 hydrogel, and fit with a second order polynomial equation. Over the validated range for each analyte, (specified in Experimental Section), the correlation coefficient was >0.9999 for all three components (Figure 3(d)-(f)). The LOQ level was established at 0.03 µg for HPMC-C_12_, 0.01 µg for PEG-b-PLA NPs, and 1 ng for Bimatoprost with sufficient sensitivity for in vitro release analysis. The method can also be generalized to hydrogels encapsulating other payload types, demonstrated by the separation of a model protein (BSA) from the hydrogel components (**Figure S2**, Supporting Information). Given the cargo Bimatoprost (LogP 3.2) represents the hydrophobicity of most small molecule therapeutics and, the C4 column is developed for analytes such as proteins and antibodies, this RPLC method should be suitable for analyzing hydrogels encapsulating a variety of synthetic and biomolecules.

**Figure 3.**
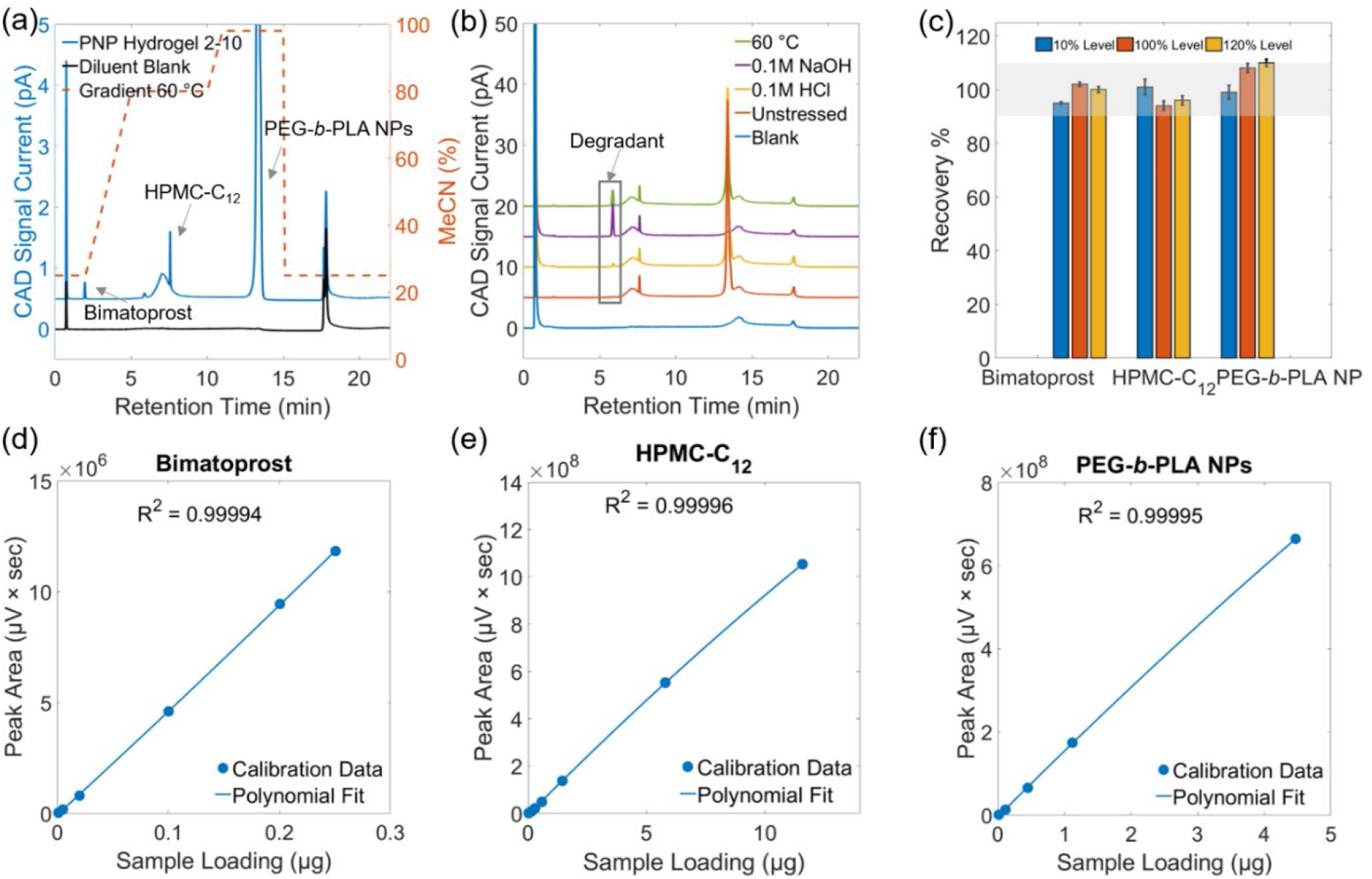
Final RPLC-CAD method chromatograms and performance. (a) RPLC-CAD chromatograms of the PNP-2-10 hydrogel (blue) and diluent blank (black), and final method gradient program (orange); (b) Forced degradation study of the HPMC-C_12_ and PEG-b-PLA; (c) Method accuracy (n=3) and precision assessment (n=3) at each concentration; Shaded area is the target range of recovery%, 90-110%. Calibration data and polynomial fitting results of (d) Bimatoprost, (e) HPMC-C_12_, and (f) PEG-b-PLA NPs.

### 2.5 Application of RPLC-CAD Method in Process Development

One application of this method was to monitor the concentrations of two polymeric components and Bimatoprost during an in vitro release study, which was included in our previous publication using a USP 7 dissolution apparatus.[25] The method was also applied here to evaluate the compatibility of E-beam sterilization with the hydrogel. E-beam sterilization is a common sterilization process for injectable formulations, involving continuous flow of high energy electrons into the treated materials.[41] However, this may lead to polymer/cargo degradation and impact rheological properties in the case of hydrogel formulations. The impact of E-beam sterilization was assessed by the RPLC-CAD method, comparing PNP hydrogels with and without E-beam treatment (Figure 4(a)). After E-beam treatment (dose range of 23-27 kGy), the Bimatoprost was found to be degraded by E-beam irradiation as evidenced by its earlier elution in the sterilized hydrogel, and its concentration was below the method’s detection limit (1 ng). Also notably, the HPMC-C_12_ profile had changed, with the peak apex eluting earlier, signifying a loss of hydrophobicity and degradation of the polymer. Oscillatory shear rheology showed altered viscoelastic properties for the hydrogel following E-beam treatment, consistent with the RPLC-CAD results suggesting gel component degradation. An amplitude sweep showed the E-beam treated hydrogel had a lower yield stress of 200 Pa compared with the untreated hydrogel’s 1500 Pa (Figure 4(b)). Hydrogel material flowed more readily in an inverted vial test following E-beam treatment (Figure 4 (b), inset). Both the angular frequency and the amplitude sweep showed the sterilized hydrogel had an order of magnitude lower moduli (Figure 4 (b) and **(c)**) and a G’ G’’ crossover point at a lower strain (Figure 4(d)), indicating the hydrogel became less stiff after E-beam irradiation. This study demonstrated the RPLC-CAD method can provide chemical stability information for the hydrogel and assess the impact of manufacturing process on the product quality. Importantly, the RPLC-CAD method can establish structure-property relationships by capturing the chemical changes and connecting those changes with rheological or other mechanical properties of the hydrogels.

**Figure 4.**
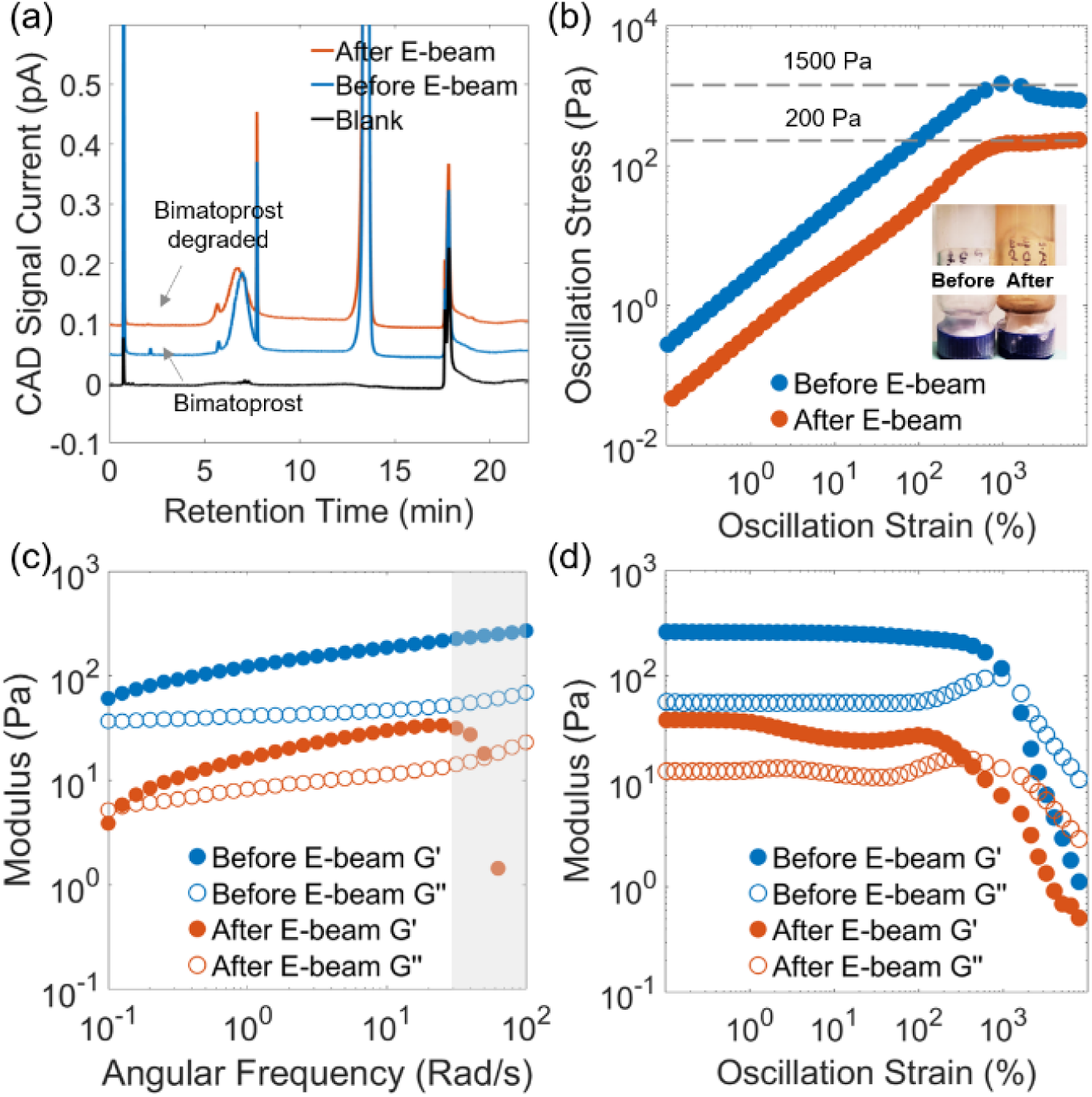
Characterization of hydrogels before and after E-beam sterilization. (a) Chemical composition analysis by the RPLC-CAD method; (b) Strain-stress characterization; Inset: photo of hydrogels before and after sterilization; Note: the color changes were limited to the vial, and the hydrogels remained as an opaque white color before and after E-beam; Moduli characterization by (c) frequency sweep and (d) amplitude sweep. The shaded data in (c) were instrument artifacts.

### 2.6 Polymer Degradant Characterization by RPLC-MS

As CAD and MS detectors both require volatile buffers as the mobile phase, the RPLC-CAD method was readily transferrable to an RPLC-MS system for higher resolution structural elucidation. Here, we probed the identity of the degradant observed in the earlier forced degradation study, particularly under the NaOH-stressed condition (Figure 3(b)). This degradant was also present at a low level in the intact hydrogel (Figure 3(a)). The mass spectrum of this degradant was collected in a qTOF mass spectrometer and processed using iFAMS.[27–31] In brief, polymer mass spectra consist of peak distributions with periodic spacing based on the mass of the repeated unit and the polymer’s net charge (Δm/z), which can be separated by Fourier transform and normalized for charge to yield much simpler mass reconstructions from multiply charged ion populations (Figure 5). The iFAMS deconvolution process, further detailed in **Figure S3** (Supporting Information), revealed that the chromatographically-observed degradant is comprised solely of 4.5-6 kDa polymer with a repeated unit of 44.05 Da, consistent with the 5 kDa PEG blocks used in the synthesis process for PEG-b-PLA copolymers.[25] Overall, the RPLC-CAD or - MS method can monitor the release of the polymeric components as well as the degradation products, such as PEG, from PEG-b-PLA, opening many opportunities to characterize and understand the molecular level structural changes during the formulation development, manufacturing, and in vivo life cycle of the supramolecular hydrogels.

**Figure 5.**
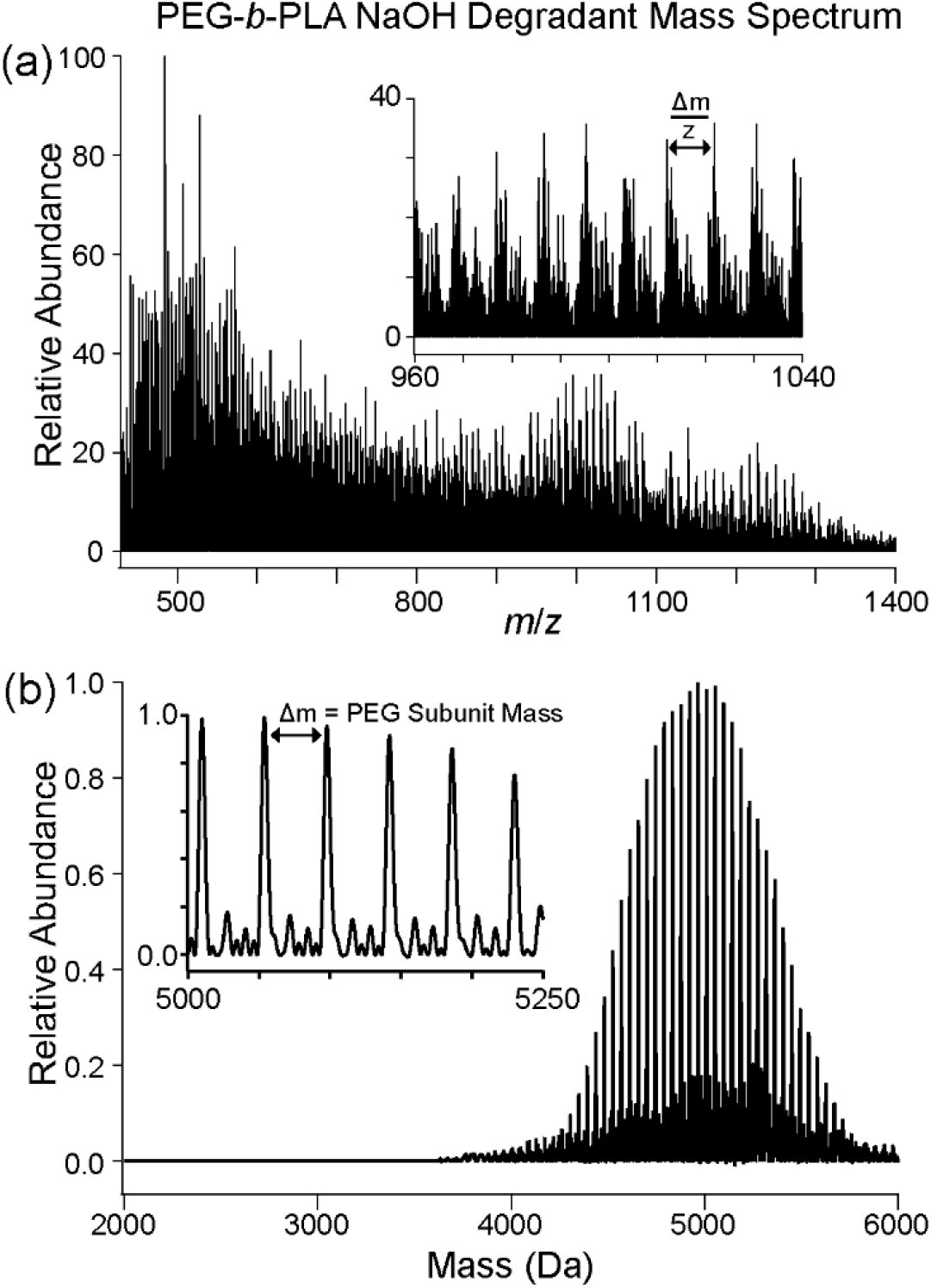
Mass spectral identification of PEG-b-PLA degradant peak from forced degradation study with NaOH. (a) 6545/XT mass spectrum of multiply charged polymers, inset indicates the periodic peak pattern observed from polymer ions. (b) iFAMS mass reconstruction of the 6545/XT spectrum, inset demonstrating the reduction in spectral complexity and identification of the repeated 44.05 Da PEG unit.

## 3. Conclusion

In summary, we presented an RPLC-CAD method as a label-free approach for characterizing a supramolecular PNP hydrogel. Instead of using the benchmark SEC separation for polymer analysis, a wide pore reversed phase column was found to provide the best specificity with a combined separation mechanism of interaction and exclusion. Coupling to a CAD detector, the RPLC-CAD method could quantitate the active cargo as well as the polymeric components with good recovery and high sensitivity. The RPLC-CAD method was applied to characterize chemical changes of the hydrogels in forced degradation and E-beam studies, showing the method could capture molecular level changes critical for the hydrogel material properties (i.e., rheological properties). In addition, the method is compatible with MS to achieve high resolution structural analysis. With assistance from the iFAMS deconvolution algorithm, the repeating unit and molecular weight of a polymer degradant were determined. Overall, this analytical method can capture the hydrogel CQAs by quantitating the composition and characterizing the polymer release, stability, and degradation of hydrogels. Based on our exploration with alternative cargos such as proteins, this analytical strategy could potentially be generalized to other supramolecular hydrogels with various cargos to obtain molecular insights and structure-property relationships for rational design and clinical translation of next generation hydrogel therapeutics.

## 4. Experimental Methods

### Materials and Reagents

USP grade HPMC, N,N-diisopropylethylamine, diethyl ether, hexanes, acetone, dimethyl sulfoxide (DMSO), acetonitrile, N-methyl-2-pyrrolidone (NMP), diazobicylcoundecene (DBU), acetic acid, formic acid, monomethoxy-PEG (5 kDa), and 1-dodecyl isocynate were purchased from Sigma-Aldrich and used as received. Lactide (LA) was purchased from Sigma-Aldrich and purified by recrystallization in ethyl acetate with sodium sulfate. Dichloromethane (DCM) was purchased from Sigma-Aldrich and further dried via cryo distillation. For HPLC analysis, deionized water was obtained from an in-house Milli-Q water filtration system. Acetonitrile (MeCN) and Tetrahydrofuran (THF) were purchased from JT Baker, and LC-MS grade trifluoroacetic acid (TFA) was purchased from Fisher Scientific. DMSO was purchased from Alfa-Aesar. Pullulan standards were purchased from Polymer Standards Service. BSA was purchased from Thermo Fisher Scientific and Bimatoprost was sourced from Toronto research chemicals.

### Synthesis of HPMC-C_12_ and PEG-b-PLA

HPMC-C_12_ and PEG-b-PLA were prepared and purified according to previously reported procedures.[25] The HPMC-C_12_ molecular weight (Mw) was determined by size exclusion chromatography coupled to multi-angle light scattering (SEC-MALS, Wyatt Technology, Santa Barbara, CA), Mw = 372 kDa (PDI=1.2). PEG-b-PLA molecular weight was determined by DMF GPC Mw= 22.5 kDa (PDI=1.07). The PEG block was 5 kDa and PLA block was 18 kDa as determined by NMR.

### Preparation of PEG-b-PLA Nanoparticles

PEG-b-PLA nanoparticles (NPs) were prepared and analyzed as previously reported.[25] A 1 mL solution of PEG-b-PLA (50 mg/mL) in 1/3 DMSO/Acetonitrile (v/v) was added dropwise to water (10 mL) (stir rate 600 rpm). Then, the NPs were purified by ultracentrifugation over a filter (molecular weight cutoff 10 kDa; Millipore Amicon Ultra-15, EMD Millipore, Burlington, MA) followed by re-suspending in Phosphate-Buffered Saline (PBS) to a final concentration of 200 mg/mL. NPs size and dispersity were characterized by dynamic light scattering (DLS, Wyatt Technology, Santa Barbara, CA) (diameter = 31.8 nm, PDI = 0.04).

### Preparation of PNP Hydrogels

PNP-2-10 hydrogels were formulated (2 wt% HPMC-C_12_ and 10 wt% PEG-b-PLA NPs) according to the previous study.[25] HPMC-C_12_ was dissolved in PBS at 6 wt% and loaded into a luer-lock syringe. A 20 wt% solution of NPs in PBS was diluted with additional PBS, containing Bimatoprost at the desired concentration, and loaded into a separate luer-lock syringe. The nanoparticle syringe was then connected to a female-female luer-lock elbow and the solution was moved into the elbow until visible at the other end. The HPMC-C_12_ syringe was then attached to the other end of the elbow with care to avoid air at the interface of HPMC-C_12_ and the NP solution. The two solutions were mixed for 1 minute or until a homogenous hydrogel was formed. After mixing, the elbow was removed and a needle of the appropriate gauge was attached.

### Instrumentation

The RPLC-CAD and SEC-CAD analysis used an Agilent 1260 series HPLC (Agilent Technologies, Santa Clara, CA) equipped with a quaternary pump, vacuum degasser, temperature controlled autosampler, thermostatted column compartment, diode array detector, and coupled to a Thermo Dionex Corona Veo RS CAD detector (Thermo Fisher Scientific, Waltham, MA). For all analysis, CAD evaporation temperature was set to 35°C, data collection was set to 5 Hz, and filter was set to 3.6 seconds.

### Chromatographic Conditions for SEC-CAD Methods

SEC separation was carried out using an Acclaim SEC-1000 7.8 × 300 mm (Thermo Fisher Scientific, Waltham, MA), a PolySep GFC-P 6000 7.8 × 300 mm (Phenomenex, Torrance, CA), and a TSKgel G5000PWXL 7.8 × 300 mm (TOSOH Bioscience, King of Prussia, PA). Mobile phase was 80% 20 mM ammonium acetate/20% MeCN (v/v) with a flow rate of 0.5-1.0 mL/min. The thermostat temperature was 30 °C. The chromatography data was processed and analyzed in Empower (Waters, Milford, MA).

### Chromatographic Conditions for RPLC-CAD Methods

RPLC separation was carried out on the following columns: Zorbax SB-CN 80 Å 3.0 × 100 mm (Agilent, Santa Clara, CA), Zorbax SB-C8 300 Å 3.0 × 150 mm (Agilent, Santa Clara, CA), Halo C4 400 Å 2.1 × 150 mm (Advanced Materials Technology, Wilmington, DE), and Halo C4 1000 Å 2.1 × 150 mm 1000 Å (Advanced Materials Technology, Wilmington, DE). Mobile phase A (MPA) was 0.05% (v/v) TFA in water and mobile phase B (MPB) was MeCN. A generic gradient was used to evaluate those columns starting at 5%MPB for 1 min, then ramping to 95%MPB over 23 mins, maintaining 95% MPB for 1 min before bringing it back to the original gradient condition. The total method run time was 30 mins. Flow rate was 0.5 mL/min for columns with 2.1 mm internal diameter (ID) and 0.8 mL/min for columns with 3.0 mm ID. Thermostat temperature was set at 30 °C unless otherwise specified.

The final optimized RPLC method used the Halo 400 Å C4 column. MPA was 0.05% TFA (v/v) in water and MPB was MeCN. Flow rate was 0.5 mL/min and column temperature was 60 °C. The sample diluent was 25% MeCN in water (v/v) unless otherwise stated. The final method gradient program was as follows: 0 – 2 min, initial hold at 25% MPB, 2 – 5 min, linear ramp from 25% to 80% MPB, 5 – 10 min, hold at 80% MPB, 10 – 11 min, linear ramp from 80% to 98% MPB, 11 – 15 min, hold at 98% MPB then the gradient was brought back to the original condition. The thermostat temperature was set at 60 °C except for E-beam experiment that conducted at 50 °C.

PEG-b-PLA STD was prepared by dissolving solid PEG-b-PLA copolymer in ACN at 1 mg/mL, then diluting with water and/or ACN to achieve the desired concentration. HPMC-C_12_ standards were prepared by adding solid HPMC-C_12_ to 25% MeCN in water (v/v), then stirring until dissolved (1-2 hrs). Hydrogel samples were dissolved using the step-wise dilution as discussed in the Diluent Study section. The chromatographic data was processed and analyzed in Empower (Waters, Milford, MA). Second order polynomial fitting was used for quantitation analysis against a multi-point calibration standard for each component.

### Chromatographic Conditions for Mass Spectrometry (MS) Analysis

For MS analysis, an Agilent 1290 series HPLC (Agilent Technologies, Santa Clara, CA) equipped with a binary pump, vacuum degasser, temperature controlled autosampler, thermostatted column compartment, and a diode array detector, was coupled to an Agilent 6545XT qTOF. Mass spectra were collected from 360-12000 m/z at a rate of 3 scans/sec. The AJS source was set at a drying temperature of 325 °C, a capillary voltage of 3000V, and a fragmentor voltage of 100V. The molecular weight and repeating unit analysis were conducted by deconvolving mass spectra from RPLC-MS total ion chromatograms with an open-source software iFAMS v.6.3 (iFAMS Quant), a Fourier-transform based algorithm developed by the Prell group to differentiate ion populations with high mass polydispersity (Figure S3, Supporting Information).[27–31]

### Method Validation

For specificity including forced degradation analysis, the HPMC-C_12_ and PEG-b-PLA NPs were stressed under acidic (0.1 M HCl, 25 °C), basic (0.1 M NaOH, 25 °C), and heated (60 °C) conditions for ∼20 hrs. The linearity was assessed over the range of 0.03 – 12 µg for HPMC-C_12_, 0.01 – 5 µg for PEG-b-PLA, and 1 – 250 ng for Bimatoprost. The linearity range was defined based on the nominal hydrogel sample composition. Accuracy and precision of each analyte was assessed at 10%, 100%, and 120% level of the nominal sample loading. The average peak response and relative standard deviation (% RSD) were calculated for each analyte at each level (n=3). Signal to noise was assessed at a sample loading of 0.03 µg for HPMC-C_12_, 0.01 µg for PEG-b-PLA, and 1 ng for Bimatoprost to determine the limit of quantitation.

## Acknowledgements

The authors acknowledged Peter Yehl, Karthik Nagapudi, Dennis Leung, Chris Crittenden, Jessie Ochoa, and Chloe Hu for their management, scientific support and expertise. E.L.M. was supported by the NIH Biotechnology Training Program (T32 GM008412). A.K.S. was supported by the National Science Foundation under award number CHE-1752994 to J.S.P. Scheme in this manuscript was created with BioRender.com

