## Supplemental Materials for "Label-Free Composition Analysis of Supramolecular Polymer – Nanoparticle Hydrogels by Reversed-Phase Liquid Chromatography Coupled with a Charged Aerosol Detector"

### 1. Supplementary Methods

*Chromatographic Conditions:*

Unless otherwise stated, all methods and conditions were the same as indicated in the main text.

The SEC-MALS-IV-dRI analysis used an Agilent 1260 series HPLC (Agilent Technologies, Santa Clara, CA) equipped with a quaternary pump, vacuum degasser, temperature controlled autosampler, thermostatted column compartment, and a diode array detector. The instrument was further coupled to the multi-angle light scattering (MALS) detector DAWN 8, inline viscometer (IV) Viscostar 3, and a differential refractometer (dRI) detector Optilab (Wyatt Technology, Santa Barbara, CA) for molecular weight (Mw), radius of gyration (Rg), and hydrodynamic radius (Rh) analysis. A TSKgel G5000PWXL 7.8×300 mm (TOSOH Bioscience, King of Prussia, PA) SEC column was used. The method mobile phase was 10 mM Potassium Phosphate (pH = 7)/Acetonitrile (80/20, v/v) with a flow rate of 0.5-1 mL/min. The thermostat temperature was 30 °C. The chromatography data was processed and analyzed in Astra 7 (Wyatt Technology, Santa Barbara, CA). The dn/dc value used for molecular weight analysis was 0.14 g/mL for HPMC-C12.[1] The dn/dc value used for PEG-b-PLA analysis was 0.043 g/mL based on the weighted average of the individual polymer block dn/dc values: PEG 0.13 g/mL and PLA 0.019.[2-3]

### 2. Supplementary Data

#### 2.1. Column Information

**Table S1.** SEC and RP Columns Used for HPMC-$C_{12}$ and PEG-*b*-PLA NP Analysis

| Column Name | Separation Mode | Phases | Particle Size (µm) | Particle Technology | Pore Size (Å) |
|---|---|---|---|---|---|
| TSKgel G5000PWXL | SEC | Hydroxylated polymethacrylate | 10 | | 1000 |
| Acclaim SEC-1000 | | Proprietary hydrophilic resin | 7 | | |
| PolySep GFC-P 6000 | | Hydrophilic polymer | Not listed | | Not listed |
| Zorbax SB-CN | RP | Cyano groups | 3.5 | Fully porous | 80 |
| Zorbax 300 Å SB-C8 | | C8 groups | 3.5 | Fully porous | 300 |
| Halo 400Å C4 | | C4 groups | 3.4 | Superficially porous | 400 |
| Halo 1000Å C4 | | C4 groups | 2.7 | Superficially porous | 1000 |

#### 2.2. Individual Polymer Characterization by SEC-MALS-IV-dRI

**Table S2.** Polymer Characterization Data by Multi-Angle Light Scattering and Inline Viscometer.

| Materials | Mw (kDa) | PDI | $R_{h,z}$ (nm) |
|---|---|---|---|
| HPMC-$C_{12}$ in 25% ACN | 372 | 1.2 | 34.8 |
| PEG-*b*-PLA STD in 25% ACN | 12521 | 1.2 | 17.2 |
| PEG-*b*-PLA NPs by nanoprecipitation in PBS | 23334 | 1.0 | 25.4 |

### 2.3 Improving HPMC-$C_{12}$ Peak Shape

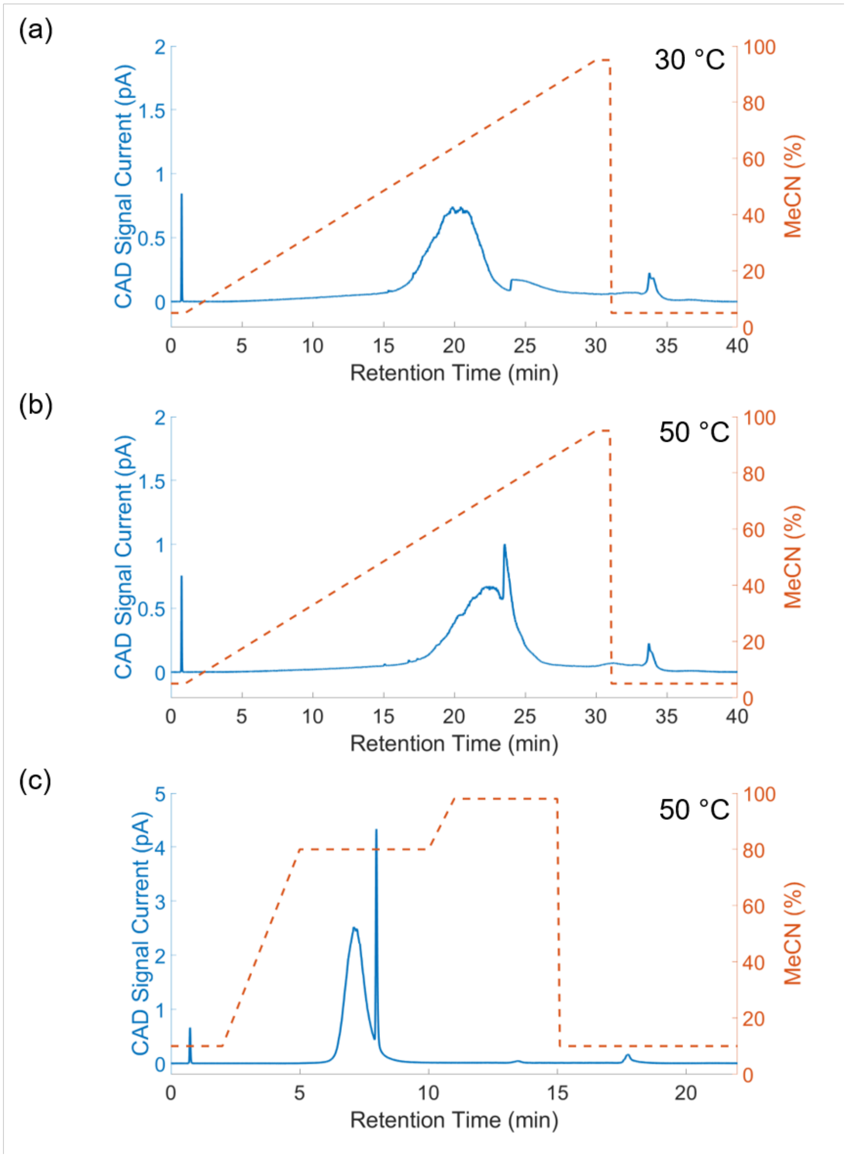


**Figure S1.** Method optimization study to improve HPMC-$C_{12}$ peak shape with a linear gradient at (a) 30 °C, (b) 50 °C, and (c) a step gradient at 50 °C.

### 2.4 Method Suitability for Alternative Cargos

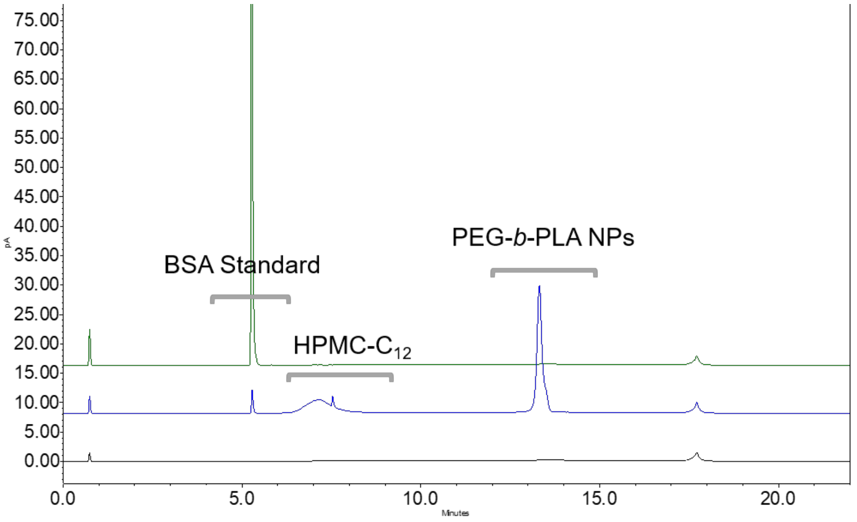


**Figure S2.** Method suitability study demonstrated by the overlay of diluent blank (black), BSA standard (green), and a mixture of BSA, HPMC-$C_{12}$ and PEG-*b*-PLA (blue), supporting its application for alternative cargos such as proteins.

### 2.5 RPLC Coupled with a High Resolution Mass Spectrometer Analysis

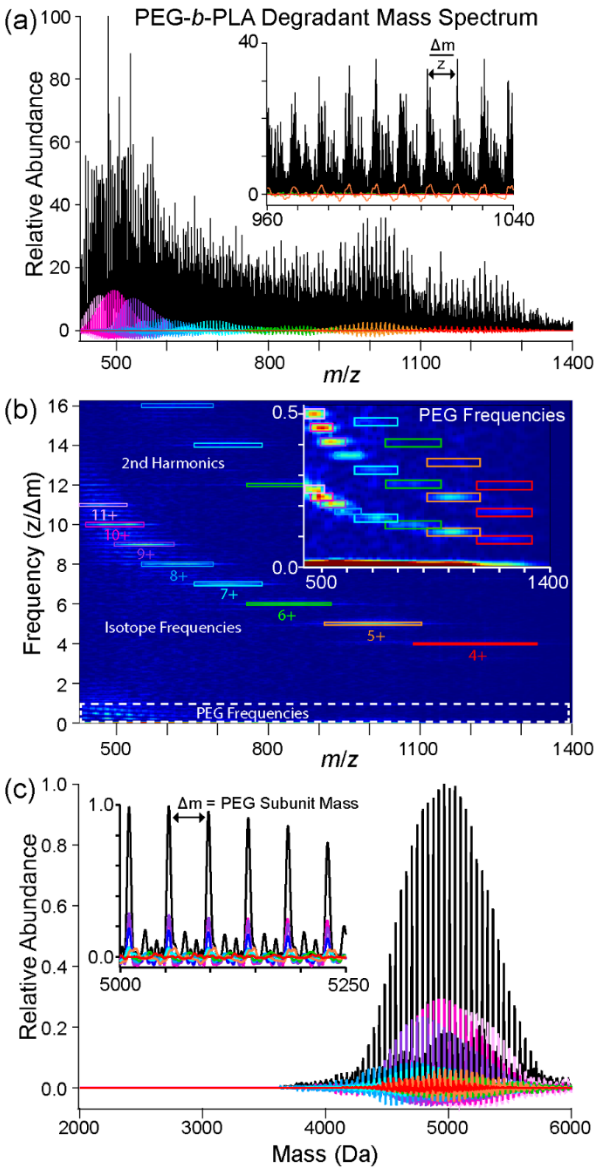


**Figure S3. iFAMS deconvolution of the PEG-*b*-PLA copolymer degradant mass spectrum using Gábor Transform (GT).**[4] Fourier analysis converts periodic peak distributions from the mass spectrum (a, inset) into distinct features in frequency (transforming Δm/z → z/Δm). GT is a windowed-Fourier transform technique that enables localization of frequency information from the mass spectrum (a) in a 2D spectrogram (b). Since the mass spectrum is isotopically resolved, GT identifies signal at integer frequencies (corresponding to isotope spacings, Δm ≈ 1 Da) in addition to lower frequencies corresponding to the PEG subunit (Δm ≈ 44.05 Da) (b, inset), simplifying charge distribution assignment of the multiply charged polymer ions. Inverse Gábor Transform of the PEG subunit frequencies and their harmonics (b, colored boxes) back to *m*/*z* generates charge-specific mass distributions with subunit resolution (a, colored spectra). The charge-specific mass distributions are normalized for charge and combined into a total mass reconstruction of the identified polymer charge series (c), and the repeated subunit mass can be confirmed from each charge state (c, inset).
